# The Effect of Genome Graph Expressiveness on the Discrepancy Between Genome Graph Distance and String Set Distance

**DOI:** 10.1101/2022.02.18.481102

**Authors:** Yutong Qiu, Carl Kingsford

## Abstract

**Motivation:** Intra-sample heterogeneity describes the phenomenon where a genomic sample contains a diverse set of genomic sequences. In practice, the true string sets in a sample are often unknown due to limitations in sequencing technology. In order to compare heterogeneous samples, genome graphs can be used to represent such sets of strings. However, a genome graph is generally able to represent a string set universe that contains multiple sets of strings in addition to the true string set. This difference between genome graphs and string sets is not well characterized. As a result, a distance metric between genome graphs may not match the distance between true string sets.

**Results:** We extend a genome graph distance metric, Graph Traversal Edit Distance (GTED) proposed by Ebrahimpour Boroojeny et al., to FGTED to model the distance between heterogeneous string sets and show that GTED and FGTED always underestimate the Earth Mover’s Edit Distance (EMED) between string sets. We introduce the notion of string set universe diameter of a genome graph. Using the diameter, we are able to upper-bound the deviation of FGTED from EMED and to improve FGTED so that it reduces the average error in empirically estimating the similarity between true string sets. On simulated TCR sequences and Hepatitis B virus genomes, we show that the diameter-corrected FGTED reduces the average deviation of the estimated distance from the true string set distances by more than 250%.

**Availability:** Data and source code for reproducing the experiments are available at: https://github.com/Kingsford-Group/gtedemedtest/

## 1 Introduction

Intra-sample heterogeneity describes the phenomenon where a genomic sample contains a diverse set of genomic sequences. A heterogeneous string set is a set of strings where each string is assigned a weight representing its abundance in the set. Computing the distance between heterogeneous string sets is essentially computing the distance between two distributions of strings. We formulate the problem of heterogeneous sample comparison as the heterogeneous string set comparison problem.

This problem can be used to compare samples where differences can be traced to the differences between sets of genomic sequences. For example, cancer samples are clustered based on differences in their genomic and transcriptomic features [1, 2] into cancer subtypes that correlate with patient survival rates. The dissimilarities between T-cell receptor (TCR) sequences are computed between individuals to study immune responses [3]. Different compositions of these sequences result in different clinical outcomes such as response to treatment.

We point out that the Earth Mover’s Distance (EMD) [4], or the Wasserstein distance [5], with edit distance as the ground metric is an elegant metric to compare a pair of heterogeneous string sets. Given two distributions of items and a cost to transform one item into another, EMD computes the total cost of transforming one distribution into another. The EMD was initially used in computer vision to compare distributions of pixel values in images [6] and later adapted to natural language processing [7]. It has also been used to approximate the distance between two genomes [8] by computing the distance between two distributions of k-mers. To compare heterogeneous string sets, when the strings and their distributions are known, we use edit distance as the cost to transform one string to another. We refer to this as the Earth Mover’s Edit Distance (EMED).

In practice, the complete strings of interest and their abundances are often unknown, and these strings are only observed as fragmented sequencing reads. It is impossible to exactly compute EMED between the true sets of complete strings from the sequencing reads only.

The challenges posed by incomplete observed sequences can be alleviated by representing the string set using a graph structure. Multiple types of genome graphs have been introduced [9, 10, 11, 12, 13, 14, 15, 16, 17, 18]. For our purposes, a genome graph is a directed multigraph with labeled nodes and weighted edges, along with a source and a sink node. A string is spelled by a source-to-sink path, or *s* − *t* path, if it is equal to the concatenation of node labels on the path. We say that a genome graph represents a string set if the union of paths that spells each string in the set is equal to the graph. In other words, a string set can be spelled by a decomposition of the genome graph.

There are several methods that compute the distance between genome graphs [19, 20, 21]. Among those, Graph Traversal Edit Distance (GTED) [21] is a general measure that can be applied to genome graphs and does not rely on the type of genome graphs nor the knowledge of the true string sets. Given two genome graphs, GTED finds an Eulerian cycle in each graph that minimizes the edit distance between the strings spelled by each cycle.

However, applying GTED on genome graphs representing heterogeneous string sets may overestimate the similarity between these string sets for two reasons. First, since GTED computes the distance between Eulerian cycles in genome graphs, it may align the prefix of a string to the suffix of another string with no additional penalties. We address this challenge by proposing an extension of GTED, called FGTED, that penalizes direct alignment of prefixes of a string with suffixes of other strings.

Second, and more significantly, both FGTED and GTED compute the edit distance between the two string sets represented by each genome graph that are most similar to each other. However, a genome graph that is constructed from sequencing fragments typically is able to represent more than one set of strings [22, 23]. As a genome graph merges shared sequences into the same node, it creates chains of bubble structures [24] that result in exponential number of possible paths, and these paths spell a much more diverse collection of strings than the original set. We call the degree to which a genome graph encodes a larger set of strings than the true underlying set the “expressiveness” of a genome graph. Due to the expressiveness of a genome graph, the Eulerian cycles found by GTED may not spell the true set of strings and the computed distance may be far from the true distance between string sets used to construct the graphs (Figure 1(a)).

**Figure 1:**
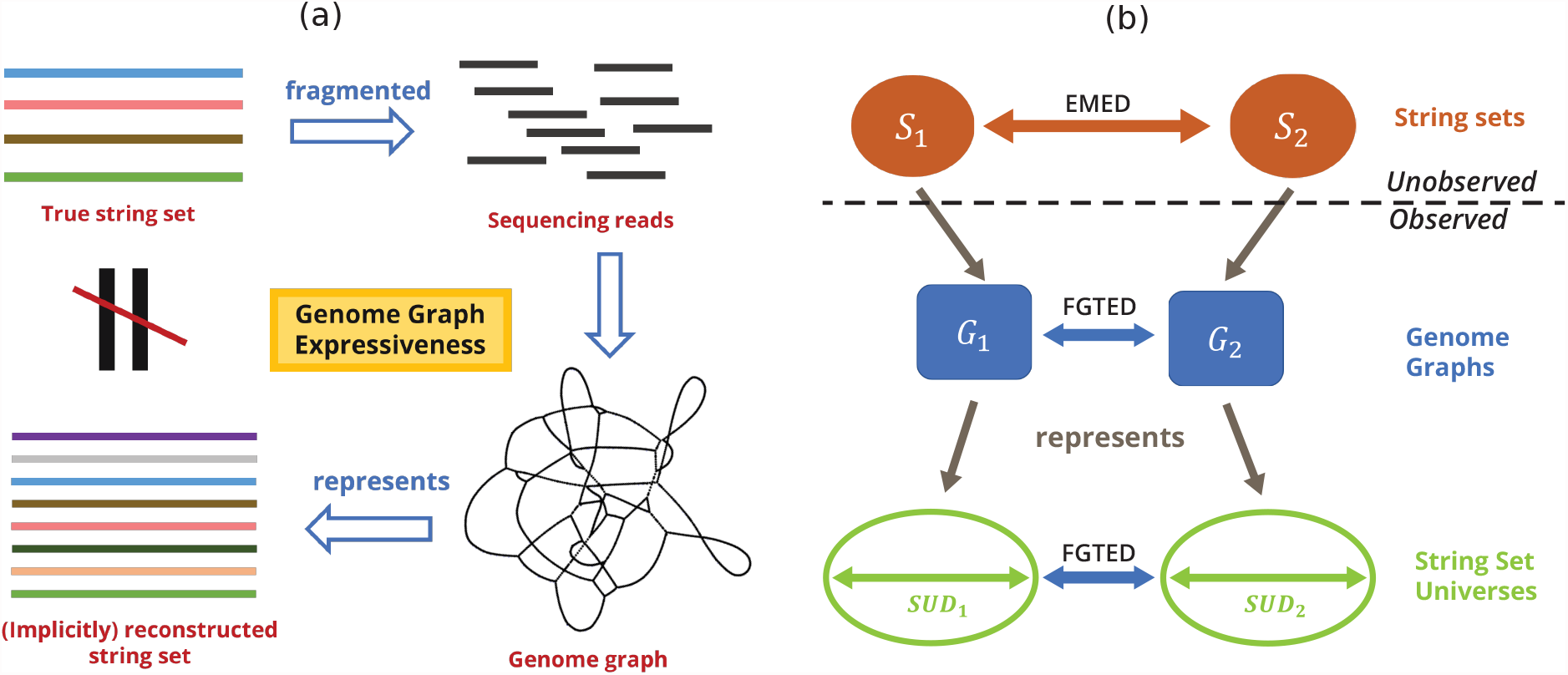
(a) Genome graph expressiveness results in inexact representations of true string sets. (b) Overview of part of theoretical contributions.

We prove both that FGTED always produces a distance that is larger than or equal to GTED, and that FGTED computes a metric that is always less than or equal to the EMED between true sets of strings.

However, FGTED and GTED can be quite far from the EMED. To resolve this discrepancy between FGTED and EMED, we define the collection of strings that can be represented by the genome graph as its string set universe, and genome graph expressiveness as the diameter of its string set universe (SUD), which is the maximum EMED between two string sets that can be represented by the graph (Figure 1(b)).

Using diameters, we are able to upper-bound the deviation of FGTED from EMED. Additionally, we are able to correct FGTED and more accurately estimate the true string set distance empirically. On simulated TCR sequences, we reduce the average deviation of FGTED from EMED by more than 300%, and increase the correlation between the true and estimated string set distances by 20%. On Hepatitis B virus genomes, we reduce the average deviation by more than 250%.

These results provide the first connection between comparisons of genome graphs that encode multiple sequences and a natural string distance and provide the first formalization of the expressiveness of genome graphs. Additionally, they provide a practical method to estimate and reduce discrepancy between genome graph distances and string set distances.

## 2 Preliminary Concepts

### 2.1 Strings

#### Definition 1 (Heterogeneous string set).

*A heterogeneous string set* 𝒮 = {(*w*_1_, *s*_1_), …, (*w*_*n*_, *s*_*n*_)} *contains a set of strings, where each string s*_*i*_ *is assigned a weight w*_*i*_ ∈ [0, 1] *that indicates the abundance of s*_*i*_ *in* 𝒮. *We say that the total weight of* 𝒮 *is* Σ_*i*∈[1,*n*]_ *w*_*i*_ = 1.

*ED*(*s*_1_, *s*_2_) is the minimum cost to transform *s*_1_ into *s*_2_ under edit distance [25]. The set of operations that transforms *s*_1_ to *s*_2_ can be written as an alignment between *s*_1_ and *s*_2_, or *A* = *align*(*s*_1_, *s*_2_). The *i*-th position in *A* is denoted by 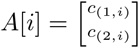, where *c*_(*a,i*)_ is either a gap character “-” or a character in *s*_*a*_.

### 2.2 Earth Mover’s Edit Distance

To find a distance between two heterogeneous string sets, we need to take into account not only the distance between pairs of strings, but also the weight, or abundance of each string in the set. When we are comparing two heterogeneous string sets, we are essentially comparing two distributions of strings. Therefore, we propose using the Earth Mover’s Distance (EMD) as a natural distance measure.

Given two distributions of items (here, strings) and a cost function that quantifies the cost of transforming one item into another, the EMD between the two distributions is the minimum cost to transform one distribution into another. Computing EMD can be viewed as a transportation problem that finds a many-to-many mapping between two sets of items and minimizes the total cost of the mapping [4, 5].

Given two heterogeneous string sets 𝒮_1_ = *{*(*w*_1_, *s*_1_), …, (*w*_*n*_, *s*_*n*_)*}* and 𝒮_2_ = *{*(*w*_*n*+1_, *s*_*n*+1_), …, (*w*_*m*_, *s*_*m*_)*}*, to compute the Earth Mover’s Edit Distance (EMED), we use the edit distance between *s*_*i*_ and *s*_*j*_ as the cost of transforming one string to another. Following procedures to compute EMD [4] as a min-cost max-flow problem, we find a mapping *M*, where *M* (*s*_*i*_, *s*_*j*_) is the amount of *s*_*i*_ ∈ 𝒮_1_ to be transformed into *s*_*j*_ ∈ 𝒮_2_, that minimizes 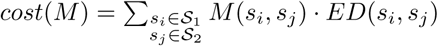. We define that *EMED*(𝒮_1_, 𝒮_2_) = min_*M*_ *cost*(*M*).

### 2.3 Flow Networks

#### Definition 2 (Valid flow network).

*A directed graph* 𝒢 = (*V, E, w*), *where w*(*e*) *is the weight of each edge, is a valid flow network if there exists a source s and sink node t such that:*

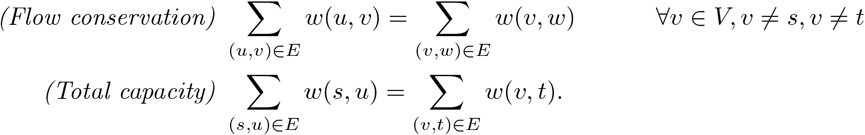

#### Definition 3 (Flow decomposition).

*A flow decomposition of a valid flow graph* 𝒢, *denoted as D*(𝒢), *is a collection of paths and their weights* 𝒫 = *{*(*w*_1_, *p*_1_), …, (*w*_*n*_, *p*_*n*_)*}, where p*_*i*_ = ((*s, u*_1_), …, (*u*_*m*_, *t*)) *is an ordered sequence of edges in* 𝒢, *such that:*

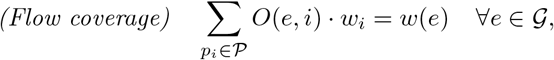

*where O*(*e, i*) *is equal to the number of occurrences of edge e in path p*_*i*_.

A valid flow network typically has more than one flow decomposition. Let the set of possible flow decompositions of 𝒢 be 𝒟_𝒢_.

### 2.4 Genome Graphs

There are many variants of genome graphs used for various purposes and in various settings. Here, we introduce the definition of genome graphs we will use.

#### Definition 4 (Genome graph).

*A genome graph* 𝒢 = (*V, E, l, w*) *is a valid flow network with node set V, edge set E, node labels l*(*u*) *for each u* ∈ *V and edge weights w*(*e*) *for each e* ∈ *E. A genome graph contains a source node s and a sink node t, and l*(*s*) =*“$”, l*(*t*) =*“#”, where* $ *and # are special characters that do not appear in any string set considered in the scope of this manuscript*.

Define operator *S*(*·*) that transforms a set of paths in a genome graph 𝒢 to a set of strings by concatenating the node labels on each path. *S*(𝒫) = *{(concat*(*p*), *w*(*p*)) | *p* ∈ 𝒫*}* is a heterogeneous string set where the weight of each string is equal to the weight of the path that spells the string.

#### Definition 5 (String set represented by a genome graph).

*A genome graph* 𝒢 *represents a string set* 𝒮 *if there exists a decomposition D*(𝒢) ∈ 𝒟_𝒢_, *such that S*(*D*(𝒢)) = 𝒮.

We use 𝒢 = *G*(𝒮) to denote when 𝒢 represents 𝒮.

#### Definition 6 (String set universe represented by a genome graph).

*The string set universe SU* (𝒢) *of a genome graph* 𝒢 *is the collection of heterogeneous string sets that can be represented by* 𝒢. *Formally, SU* (𝒢) = *{S*(*D*) | *D* ∈ 𝒟_𝒢_ *}*.

### 2.5 Alignment Graph

An alignment graph is used to align two genome graphs [21] and can be viewed as a graph product between two genome graphs. A special case of the alignment graph [26] is used to align a string to a graph where the string is represented as a graph with only one path. We assume that the genome graphs to be aligned are transformed so that the label of each node contains only one character.

#### Definition 7 (Alignment graph).

*Given genome graphs* 𝒢_1_ = (*V*_1_, *E*_1_, *l*_1_, *w*_1_) *and* 𝒢_2_ = (*V*_2_, *E*_2_, *l*_2_, *w*_2_), *an alignment graph AG*(𝒢_1_, 𝒢_2_) = (*V*_*A*_, *E*_*A*_, *cost, w*) *is a directed graph with node set V*_*A*_, *edge set E*_*A*_, *edge cost cost*(*e*) *and edge weight w*(*e*) *for each edge e* ∈ *E*_*A*_. *The alignment graph is defined following the steps:*

- *V*_*A*_ *is constructed by adding pairings of nodes in V*_1_ *and V*_2_; *that is V*_*A*_ = *{*(*u*_1_, *u*_2_) | *u*_1_ ∈ *V*_1_, *u*_2_ ∈ *V*_2_*}*.
- *For each edge* (*u*_1_, *v*_1_) ∈ *E*_1_ *and* (*u*_2_, *v*_2_) ∈ *E*_2_, *create three types of edges:*
  1. *A match/mismatch edge e* = ((*u*_1_, *u*_2_), (*v*_1_, *v*_2_)) *with w*(*e*) = min*{w*_1_(*u*_1_, *v*_1_), *w*_2_(*u*_2_, *v*_2_)*}*.
  2. *An insertion (in) edge e* = ((*u*_1_, *u*_2_), (*u*_1_, *v*_2_)) *with w*(*e*) = *w*_2_(*u*_2_, *v*_2_).
  3. *A deletion (del) edge e* = ((*u*_1_, *u*_2_), (*v*_1_, *u*_2_)) *with w*(*e*) = *w*_1_(*u*_1_, *v*_1_).

*The cost of an in/del edge and a mismatch edge is equal to a customized penalty. The cost of a match edge is equal to zero. A match/mismatch edge should be distinguished with an in/del edge if the corresponding edge in one of the input graphs is a self-loop*.

Each edge *e* = ((*u*_1_, *u*_2_), (*v*_1_, *v*_2_)) in an alignment graph can be projected onto one edge in each of the input graphs. An edge in each of the input graphs can also be projected onto a set of edges in *AG*.

#### Definition 8 (Projection function).

*Define the projection function as P*_(𝒢, ℋ)_(*e*) = *E*′ *that maps an edge e from graph* 𝒢 *to a set of edges E*′ *in graph* ℋ. *The projection function maps an edge in the alignment graph to the edges in the input graphs that are matched together by that edge. It also maps an edge in one of the input graphs to a set of edges in the alignment graph where it is matched with other edges in another input graph. Specifically:*

*Projection from alignment graph to one of the input graphs is defined by*

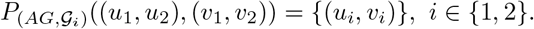

*Projection from one of the input graphs to alignment graph is defined by*

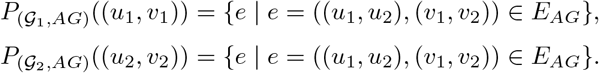

Given a set of paths 𝒫 in *AG*, we use 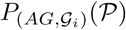 to denote the projection of 𝒫 onto 𝒢_*i*_, where 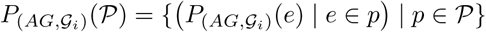.

For convenience, we define that 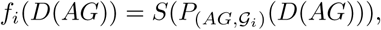, which is the set of strings spelled by a path decomposition in *AG* that is projected onto 𝒢_*i*_.

### 2.6 Graph Traversal Edit Distance (GTED)

Graph Traversal Edit Distance (GTED), proposed by Ebrahimpour Boroojeny et al. [21], is a distance between two labeled graphs which are assumed to be Eulerian graphs. Given a genome graph in our definition, we add an edge directing from sink to source with weight equal the sum of edge weights that are directing from the source node in order to make an Eulerian graph.

Let the language of 𝒢, *L*(𝒢), be the set of strings spelled by Eulerian cycles in 𝒢. Formally, *L*(𝒢) = *{S*(*c*) | *c* is an Eulerian cycle in 𝒢*}*.

#### Definition 9 (Graph Traversal Edit Distance [21]).

*Let* 𝒢_1_ *and* 𝒢_2_ *be two Eulerian graphs, where the weights on the edges are seen as the number of times an edge must be visited in each Eulerian cycle. Then*,

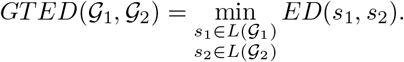

GTED finds one Eulerian cycle in each genome graph such that the edit distance between the strings spelled by the Eulerian cycles is minimized. GTED is computed by solving a linear programming (LP) formulation (Equations (1)-(4)) on the alignment graph *AG*(𝒢_1_, 𝒢_2_). The LP formulation is as follows:

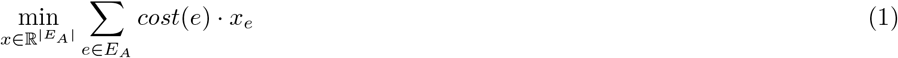

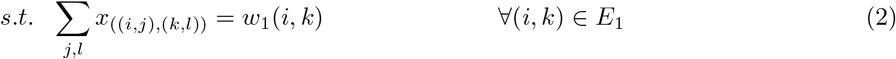

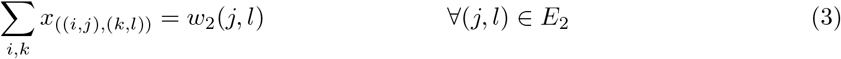

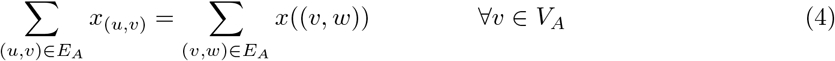

Ebrahimpour Boroojeny et al. [21] prove that GTED is equal to the optimal solution of this LP formulation.

## 3 An Extension of GTED

GTED was originally used to compare genome graphs that are assumed to contain single genomes. It is therefore intuitive that each string represented by the genome graph is spelled with an Eulerian cycle. This property follows the property of assembly graphs [27]. When the genome graph represents more than one string, finding a string spelled by an Eulerian cycle *c* in the graph is equivalent to finding a concatenation of a permutation of strings in a string set. When aligning two Eulerian cycles, *c*_1_ and *c*_2_, from input graphs, the boundaries between strings are ignored and the prefix of one string may be aligned to the suffix of another string with no cost. However, such alignment is not allowed when we align sets of strings using EMED.

We propose an extension of GTED with a modified cost function in edit distance computation so that the cost of aligning the sink character # with any other character is infinity.

Figure 2(a) shows an example of the alignment graph built from two input graphs using the proposed cost function. Let the sink nodes in 𝒢_1_ and 𝒢_2_ be *t*_1_ and *t*_2_, and the source nodes be *s*_1_ and *s*_2_, respectively. After removing all the alignment edges with infinite costs, there is an edge to the alignment node (*t*_1_, *t*_2_) in *AG* if and only if there exists an edge (*u*_1_, *t*_1_) in 𝒢_1_ and an edge (*u*_2_, *t*_2_) in 𝒢_2_. The only incoming edge to (*s*_1_, *s*_2_) is ((*t*_1_, *t*_2_), (*s*_1_, *s*_2_)). We refer to the edge ((*t*_1_, *t*_2_), (*s*_1_, *s*_2_)) as the sink-to-source edge, or *t* − *s* edge in alignment graph in the rest of the manuscript.

**Figure 2:**
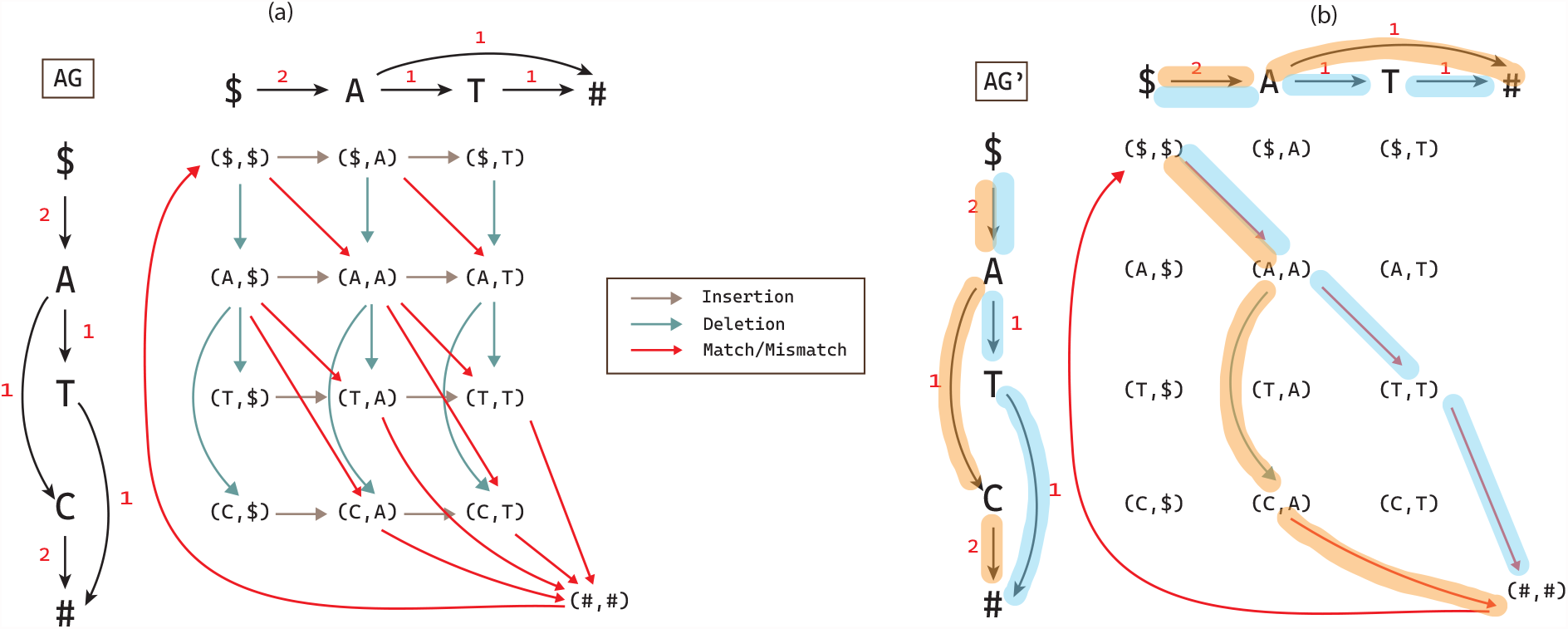
(a) An alignment graph *AG* between 𝒢_1_ (vertical) and 𝒢_2_ (horizontal). Insertion, deletion and match/mismatch edges are labeled with different colors. (b) *AG*\* after removing all the edges with zero flow in a solution to *FGTED*(𝒢_1_, 𝒢_2_). Edges in 𝒢_1_ and 𝒢_2_ that are highlighted with matching colors are projections from edges in *AG*\* to 𝒢_1_ and 𝒢_2_, respectively. Path ($,*A, T*, #) ∈ 𝒢_1_ is aligned to ($,*A, T*, #) ∈ 𝒢_2_ and path ($,*A, C*, #) ∈ 𝒢_1_ is aligned to ($,*A*, #) ∈ 𝒢_2_. The weights on *AG* and *AG*\* edges are omitted for simplicity.

We let Flow-GTED, or FGTED, denote the distance computed using the alignment graph after removing all infinity cost edges that forbid aligning the sink with any nodes other than the source node.

### Theorem 1.

*GTED*(𝒢_1_, 𝒢_2_) ≤ *FGTED*(𝒢_1_, 𝒢_2_) *for any pair of genome graphs* 𝒢_1_, 𝒢_2_.

*Proof*. Since FGTED is computed on a smaller alignment graph that contains fewer edges than that for computing GTED, FGTED explores a smaller solution space than GTED in solving the LP formulation. Therefore, any feasible solution to the LP formulation for *FGTED*(𝒢_1_, 𝒢_2_) is a feasible solution to the LP formulation for *GTED*(𝒢_1_, 𝒢_2_). Since *GTED*(𝒢_1_, 𝒢_2_) minimizes the objective, the theorem is true. □

## 4 The Relationship Between GTED, FGTED and EMED

Let *AG*\* be the alignment graph after removing the *t* − *s* edge and all the edges from *{e* | *x*_*e*_ = 0, *e* ∈ *E*_*A*_*}* from the LP solution to Equations (1)–(4). We say that *AG*\* is a solution of FGTED. Due to constraints (2)–(4), *AG*\* is a valid flow network. Let *D*(*AG*\*) be a flow decomposition in *AG*\*. Similar to the Eulerian cycles found during the GTED computation, each path in *D*(*AG*\*) can be projected to a path in 𝒢_1_ and a path in 𝒢_2_.

Denote 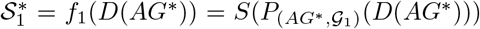 as the set of strings spelled by the set of projected paths from a decomposition *D*(*AG*\*) to 𝒢_1_. Similarly, 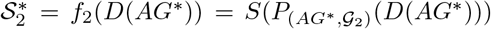. We show an example of path projections in Figure 2(b).

Observing that we can do flow decomposition in both the FGTED solution and input genome graphs, we will show in this section that FGTED can be bounded by EMED between decompositions in input genome graphs and in the alignment graph solutions.

### Theorem 2.

*Given two sets of strings* 𝒮_1_ *and* 𝒮_2_, *and genome graphs representing these string sets*, 𝒢_1_ = *G*(𝒮_1_) *and* 𝒢_2_ = *G*(𝒮_2_),

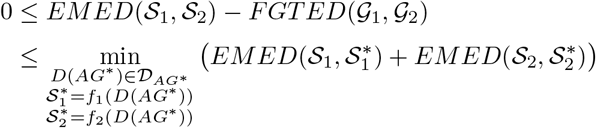

*where AG*\* *is the solution obtained from FGTED*(𝒢_1_, 𝒢_2_).

The proof of this theorem is completed in two parts. The first inequality is proven in Section 4.1 and the second is proven in Section 4.2. Since FGTED computes a distance that is larger than GTED between the same pair of genome graphs (Theorem 1), Theorem 2 also shows that FGTED always estimates the distance between true string sets more accurately than GTED.

### 4.1 FGTED is Always Less Than or Equal to EMED

We show in this section that FGTED can be expressed in terms of EMED between string sets constructed from decomposing *AG*\*. In other words, similar to GTED, FGTED finds a decomposition in 𝒢_1_ and 𝒢_2_, respectively, that minimizes the EMED between them. Analogous to Definition 9, we have:

#### Theorem 3.

*Given two genome graphs* 𝒢_1_ *and* 𝒢_2_,

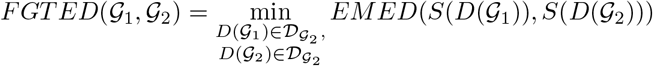

To prove Theorem 3, we first explore the relationship between an *s* − *t* path in *AG*\* and the strings spelled by the projections of this path onto 𝒢_1_ and 𝒢_2_.

#### Lemma 1.

*Given an s-t path p* ∈ *D*(*AG*\*), *let* 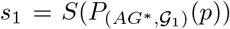 *be the string spelled by projecting p onto* 𝒢_1_, *and* 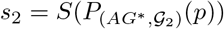. *Then for any p* ∈ *D*(*AG*\*),

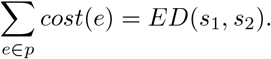

*Proof*. We prove in two directions.

(≥ **direction)** We construct *A* = *align*(*s*_1_, *s*_2_) from *p*. For each *e* = ((*u*_1_, *u*_2_), (*v*_1_, *v*_2_)) ∈ *p*: (1) if *u*_1_ = *v*_1_, add 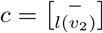 to *A*, (2) if *u*_2_ = *v*_2_, add 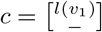 to *A*, (3) else, add 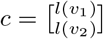 to *A*. By definition of an alignment graph, *cost*(*e*) = *cost*(*c*) in for all *e*, and therefore *cost*(*A*) = Σ_*c*∈*A*_ *cost*(*c*) = Σ_*e*∈*p*_ *cost*(*e*). Since edit distance minimizes the cost of edit operations, *cost*(*A*) = *cost*(*p*) ≥ *ED*(*s*_1_, *s*_2_).

(≤ **direction)** We construct *p*′ from *A*\* = *align*(*s*_1_, *s*_2_) such that *cost*(*A*\*) = *ED*(*s*_1_, *s*_2_). The procedure is similar as above — for each pair of adjacent entries in *A*\*, add corresponding edge to *p*′. Then *cost*(*p*′) = *cost*(*A*\*) = *ED*(*s*_1_, *s*_2_).

Let *AG*′ = *AG*\* *\ p* ∪ *p*′. Both *p* and *p*′ can be found in *AG*, and both *p* and *p*′ can be constructed by the alignment of the same pair of strings. Therefore, *AG*′ is also a valid flow network and a feasible solution to FGTED. Since *AG*\* is the optimal solution to *FGTED, cost*(*AG*\*) ≤ *cost*(*AG*′), and

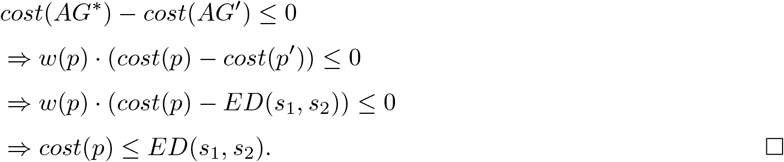

We have shown that the cost of an *s* − *t* path in *AG*\* is equal to the edit distance between its projections onto input graphs. Using this lemma, we can transform an optimal FGTED solution into an EMED solution.

Given an optimal FGTED solution, *AG*\*, let the set of possible flow decompositions of *AG*\* be 𝒟_*AG*\*_. Let *D*(*AG*\*) be one of the flow decompositions that is a set of weighted *s* − *t* paths. We can construct heterogeneous string sets 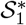 and 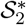 by projecting paths in *D*(*AG*\*) to 𝒢_1_ and 𝒢_2_. Formally, 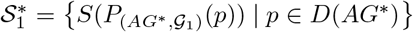 and 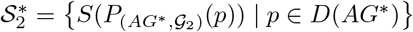.

#### Lemma 2.

*Given* 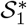 *and* 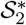 *obtained from any decomposition D*(*AG*\*) ∈ 𝒟_*AG*\*_,

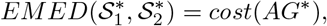

*where cost*(*AG*\*) *is sum of edge costs in the solution alignment graph to FGTED*.

*Proof*. We prove in two directions.

(≤ **direction)** We construct a mapping *M* between strings in 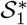 and 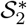 from the decomposition *D*(*AG*\*), where *M* (*s*_*i*_, *s*_*j*_) is the portion of *s*_*i*_ ∈ 𝒮_1_ and *s*_*j*_ ∈ 𝒮_2_ that are aligned. For each *p* ∈ *D*(*AG*\*), we obtain *s*_1_ and *s*_2_ as strings constructed from projections of *p* onto 𝒢_1_ and 𝒢_2_ and increment the weight of mapping *M* (*s*_1_, *s*_2_) by *w*(*p*). After iterating through all paths in *D*(*AG*\*), the cost of *M* is

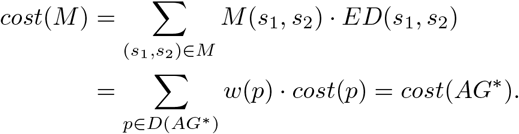

*M* is also a feasible solution to the LP formulation of EMED. Since *EMED* minimizes the cost of mapping between 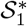 and 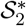, 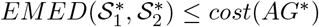.

(≥ **direction)** We construct a valid flow network, *AG*′ using an optimal solution to 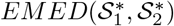. For each pairing (*s*_*i*_, *s*_*j*_) for *s*_*i*_ ∈ 𝒮_1_ and *s*_*j*_ ∈ 𝒮_2_, we obtain its weight *w* and cost *c* from the EMED solution. Let *A* = *align*(*s*_*i*_, *s*_*j*_) be an optimal alignment under edit distance, and *cost*(*A*) = *c*. We then add a path corresponding to *A* with weight *w* in *AG*′. This follows the same procedure in the proof of Lemma 1. After adding all paths, we obtain *AG*′ with 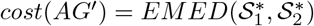. Since *cost*(*AG*\*) is minimized by FGTED, 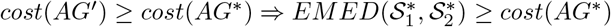. □

Lemma 2 provides a transformation algorithm between optimal solutions to EMED and solutions to FGTED. Using Lemma 2, we can show that the EMED between 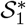 and 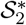 constructed from any decomposition in *AG*\* is equal to the decompositions of 𝒢_1_ and 𝒢_2_ that are closest in terms of EMED.

#### Lemma 3.

*Given* 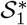 *and* 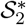 *obtained from any decomposition D*(*AG*\*) ∈ 𝒟_*AG*\*_,

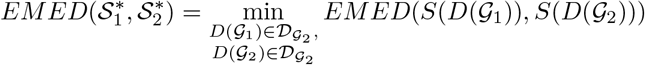

*Proof*. In Lemma 2, 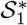 and 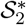 can be constructed from decomposing 𝒢_1_ and 𝒢_2_. Suppose for contradiction that there exists a decomposition that constructs string sets 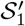 and 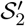, such that 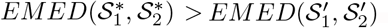. Following the procedure in the proof of Lemma 2, we can construct a feasible solution to FGTED with cost equal to 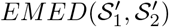, which is less than 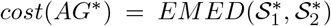. This contradicts with the assumption that FGTED minimizes *cost*(*AG*\*). □

Theorem 3 is therefore true because of Lemma 2 and 3. Using Theorem 3, we are able to prove the first inequality in Theorem 2 with Lemma 4.

#### Lemma 4.

*Given heterogeneous string sets* 𝒮_1_ *and* 𝒮_2_ *and genome graphs representing these string sets*, 𝒢_1_ = *G*(𝒮_1_) *and* 𝒢_2_ = *G*(𝒮_2_), *FGTED*(𝒢_1_, 𝒢_2_) ≤ *EMED*(𝒮_1_, 𝒮_2_).

*Proof*. Given Theorem 3, FGTED finds flow decomposition in 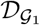 and 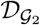 that minimizes the EMED between them. Since 𝒮_1_ and 𝒮_2_ can be constructed from a flow decomposition in 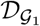 and 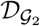, respectively, this lemma is true. □

### 4.2 Genome Graph Expressiveness

A genome graph typically can represent more than one set of strings. We name the collection of string sets representable by a genome graph the *string set universe* of that genome graph, or *SU* (𝒢). Using Theorem 3, we can say that FGTED finds two sets of strings in the string set universe of 𝒢 that are closest in the metric space of EMED. We define the expressiveness of a genome graph as the diameter of its string set universe, which is the maximum EMED between the string sets in *SU* (𝒢).

#### Definition 10 (String Set Universe Diameter (SUD)).

*Given a genome graph* 𝒢,

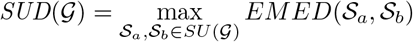

#### 4.2.1 String Set Universe Diameter as an Upper Bound on Deviation of FGTED from EMED

The string set universe diameter gives one measure of the size of *SU* (*G*), and it can also be used to characterize the deviation of GTED from EMED.

Recall that 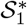 and 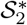 are string sets obtained from a decomposition *D*(*AG*\*), and that 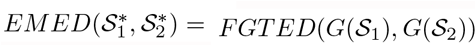, where 𝒮_1_ and 𝒮_2_ are true string sets. From Theorem 2, we have that 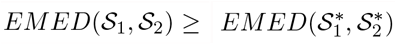. We can bound the deviation of 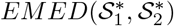 from *EMED*(𝒮_1_, 𝒮_2_) using triangle inequalities.

##### Lemma 5.

*Given string sets* 𝒮_1_ *and* 𝒮_2_ *and genome graphs* 𝒢_1_ = *G*(𝒮_1_) *and* 𝒢_2_ = *G*(𝒮_2_),

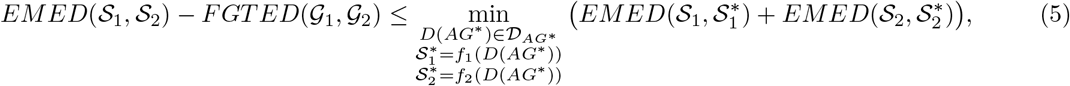

*where AG*\* *is the solution obtained from FGTED*(𝒢_1_, 𝒢_2_).

*Proof*. Both edit distance and EMD are metrics [4, 25], which means that triangle inequality holds for EMED between strings. Therefore, for any string sets 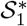 and 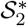,

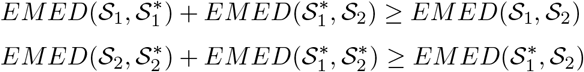

Combining two inequalities, we have

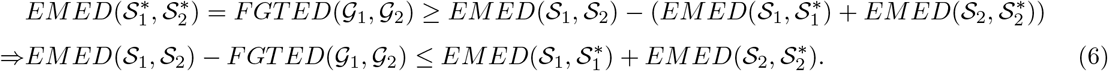

The above inequality (6) holds for any string sets 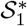 and 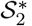;. To give a tight upper bound on the deviation, we take the minimum over all possible pairs of string sets constructed from decomposing *AG*\* that yields inequality (5). □

Lemma 5 proves the second inequality of Theorem 2 thus completing the proof for Theorem 2 with Lemma 4.

The upper-bound found in Lemma 5 can be used as a factor that evaluates the pair-wise expressiveness of two genome graphs. While a genome graph may represent a large universe of string sets, as long as the true string set is close to the “best” string set in the pair-wise comparison, the deviation of FGTED from EMED is small. We define this upper bound as the String Universe Co-Expansion Factor (SUCEF), which can be used to evaluate the discrepancy between FGTED and EMED.

##### Definition 11 (String Universe Co-Expansion Factor (SUCEF)).

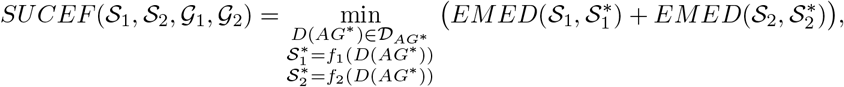

*where AG*\* *is the solution to FGTED*(𝒢_1_, 𝒢_2_).

On the other hand, finding SUCEF not only requires knowledge of true string sets 𝒮_1_ and 𝒮_2_, but SUCEF is also a pair-dependent measure that needs to be calculated for every pair of string sets and corresponding genome graphs. In order to characterize the effect of the expressiveness of individual genome graphs, we derive another upper bound on the deviation of FGTED from EMED using the string set universe diameters.

The sum of string set universe diameters of two genome graphs is an upper bound on SUCEF of these graphs and any two sets of strings they represent.

##### Lemma 6.

*Given two genome graphs* 𝒢_1_ *and* 𝒢_2_ *and two sets of strings* 𝒮_1_ *and* 𝒮_2_ *they represent*,

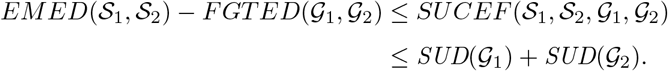

*Proof*. Both 𝒮_1_ and 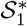 are represented by 𝒢_1_ and belong to *SU* (𝒢_1_). Therefore, by definition of string set universe diameter, 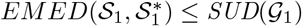 as the diameter maximizes the distance between any pair of strings represented by the genome graph. The same holds for 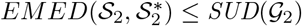. □

Using Lemma 6, we can now bound the deviation of FGTED from EMED using the expressiveness of individual genome graphs. In the following sections, we show that we can empirically estimate the anticipated discrepancy between FGTED and EMED using string set universe diameters, and we can produce improved FGTED which reduces the anticipated deviation from EMED and better correlates with EMED.

## 5 Practically Correcting the Discrepancy between FGTED and EMED

### 5.1 Estimating String Set Universe Diameters

The string set universe diameter of a genome graph can be estimated by sampling flow decompositions of the graph. To sample a flow decomposition, we first sample one *s* − *t* path. At each node *u*, we choose the neighbor *v* with the highest edge weight *w*(*u, v*) with probability 0.5 and randomly choose a neighbor otherwise. After sampling a path, we send flow that is equal to the minimum edge weight on that path and produce the residual graph by subtracting the flow from edge weights on that *s* − *t* path. We repeat this process on the residual graph until all edge weights are zero. This process assumes that the input genome graphs are acyclic to ensure all edge capacities (weights) are satisfied. If a genome graph is cyclic, string sets from *SU* (𝒢_1_) can be obtained by sampling Eulerian cycles in the genome graph. After sampling 50 pairs of flow decompositions, we construct string sets from sampled flow decompositions and calculate pairwise EMED. We then obtain the highest pairwise EMED and use it as the estimated diameter.

### 5.2 Correcting FGTED Using String Set Universe Diameters

Using sum of SUDs, we empirically estimate the deviation of FGTED from EMED with a linear regression model. We denote the deviation of FGTED from EMED by *deviation*(𝒮_1_, 𝒮_2_, 𝒢_1_, 𝒢_2_), which is computed as |*EMED*(𝒮_1_, 𝒮_2_) − *FGTED*(*G*(𝒮_1_), *G*(𝒮_2_))|. The linear regression model, *LR*, has the following form

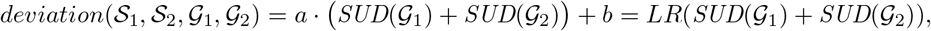

where *a* is the coefficient of the model and *b* is the intercept. The fitted model will minimize the mean squared error between predicted deviation and true deviation in the training set.

The corrected FGTED for each pair of graphs is calculated using the learned linear regression model as follows.

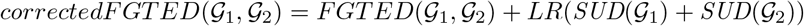

The deviation of corrected FGTED from EMED has the same form as the deviation of uncorrected FGTED from EMED.

### 5.3 Data

We evaluate the use of string set universe diameters on two sequence sets:

1. **Simulated T-Cell Receptor (TCR) Repertoire**. We simulate 50 sets of TCR sequences and assign weights to each sequence using reference gene sequences of V, D and J genes from Immunogenetics (IMGT) V-Quest sequence directory [28]. The number of sequences in each set varies from 2 to 5. We then generate 225 pairs of TCR string sets. Each TCR sequence is about 300 base pairs long. See the Appendix for detailed simulation process.
2. **Hepatitis B Virus (HBV) Genomes**. We collect 9 sets of HBV genomes from three hosts — humans, bats and ducks — from the NCBI virus database [29]. We build 36 pairs of HBV string sets. See the Appendix for detailed string set construction process.

We construct a partial order MSA graph on each string set [30]. We first conduct multiple sequence alignment (MSA) for each string set using Clustal Omega [31]. Then for each column of the MSA, we create a node for each unique character and add an edge between two nodes if the characters in node labels are adjacent in the input strings at that column. For each consecutive stretch of gap characters, no nodes are created, but an edge is added between flanking columns of the stretch of gaps. We also create a source node and a sink node that are connected to nodes representing the first and last characters of the input strings. The MSA graphs created in this process are all acyclic. We compute FGTED on MSA graphs by adding sink-to-source edges.

### 5.4 Corrected FGTED More Accurately Estimates Distance Between Unseen String Sets Encoded With Genome Graphs

We compute EMED and FGTED on string set pairs and genome graph pairs. Computing FGTED takes about 1 hour using 10 threads for each MSA graph of the simulated TCR sequence sets and 4 hours on using 10 threads for each MSA graph on the HBV genomes. The LP optimizations are done using the Gurobi solver [32].

For each pair of string sets, we obtain the deviation of FGTED from EMED and sum of estimated SUDs. We fit two linear regression models, *LR*_*TCR*_ and *LR*_*HBV*_, to predict deviation from sum of SUDs on simulated TCR sequences and HBV genomes separately.

We evaluate the corrected and uncorrected FGTED by performing Pearson correlation experiments. We fit *LR* models on half of the data and compute the corrected FGTED on the other half as the test set. We evaluate the correlation between corrected and uncorrected FGTED and EMED on the test set. Two-tail P-values are calculated for each correlation experiment to test for non-correlation.

The *LR* models are evaluated with 10-fold cross validation. We randomly permute and split data into 10 equal parts. In each of the 10 iterations, we use one part as the test set and the rest as the training set. An average deviation is calculated across all iterations.

In Table 1, we show that using string set universe diameters, we are able to improve the correlation between FGTED and EMED on both the simulated TCR sequences and HBV genomes (Figure 3). Both Pearson correlation experiments are statistically significant with P-values *<* 0.01. On HBV genomes, since the correlation between uncorrected FGTED and EMED is approaching 1, no significant improvement is observed. On the other hand, significant reduction in average deviation is observed on both types of data. We are able to reduce the average deviation from 32.74 to 9.13 on simulated TCR sequences and from 140.12 to 54.87 on HBV genomes.

**Table 1:** Pearson correlation and average deviation between EMED and corrected and uncorrected FGTED on simulated TCR and HBV sequences. Pearson correlation is calculated on a held-out set of data for both simulated TCR and HBV that consist of 50% of data, and *LR* model is fit on the other half. Average deviation is calculated by taking the average of mean deviations across 10-fold cross validation.

| FGTED | Pearson Correlation |  | Average Deviation |  |
| --- | --- | --- | --- | --- |
|  | Simulated TCR | HBV | Simulated TCR | HBV |
| Uncorrected | 0.74 | **0.99** | 32.74 | 140.12 |
| Corrected | **0.90** | **0.99** | **9.13** | **54.87** |

**Figure 3:**
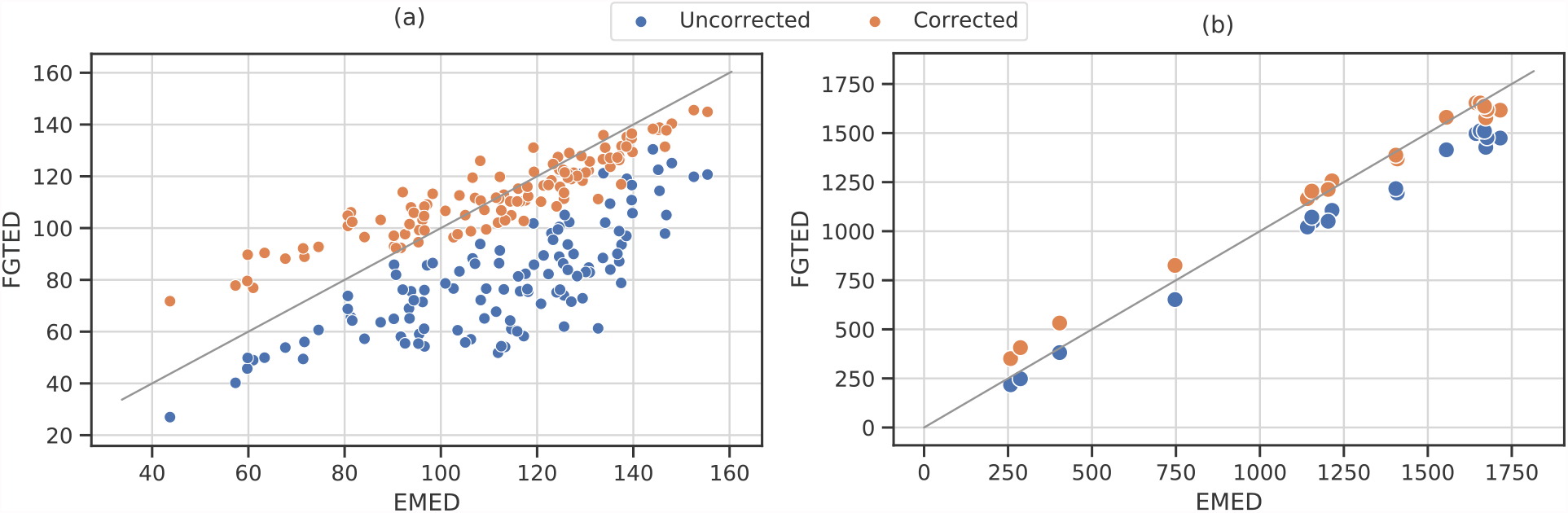
Correlation between EMED and FGTED before and after correction with string set universe diameters on half of (a) simulated TCR sequences and (b) HBV genomes. The relationship between EMED and uncorrected FGTED is shown in blue and corrected FGTED is shown in orange. Grey lines signify equality between EMED and FGTED. The data shown in Figure 3 are held-out sets from TCR and HBV genomes that consist of 50% of the data. The corrected FGTED is computed using the LR models fit on the other half.

One caveat of using SUDs for correcting distances between genome graphs is that this correction is not guaranteed to always improve the distance. Given two string sets, there is always an adversarial worst case where adjusting the distance using this approach reduces the accuracy in estimating string sets distances. As seen from Figure 3, when EMED between true string sets are small, the corrected FGTED may overestimate the EMED and result in a larger deviation. Nevertheless, we show that corrected FGTED reduces the anticipated deviation from EMED.

## 6 Discussion

A genome graph’s string set universe diameter (SUD) provides information on the size and diversity of the represented string sets. We show that we can use SUDs to practically characterize the discrepancy between GTED and EMED and to obtain a more accurate distance between unseen string sets encoded in genome graphs on average. While the results are obtained on short genomic sequences due to the high computational cost of FGTED and GTED, this result is encouraging, and string set universe diameters could be also used on approximate genome graph comparison methods to evaluate and improve their accuracies [19, 20].

SUDs could also be used to characterize the diversity of strings represented by reference genome graphs that are used in sequence-to-graph alignment [33, 34]. In sequence-to-graph alignment, it is often desired that a more diverse set of strings than the original reference string set is represented by the graph. Here, SUDs could be used as a measure to control the right amount of variation in the string set universe of created genome graphs.

Another future direction is to use expressiveness as a regularization term in the objective function to construct better genome graphs. To ensure efficiency of genome graphs in storing sequences, we can construct genome graphs that minimize their sizes [35, 36]. However, only optimizing for the size of a genome graph may result in graphs that are highly expressive, and the distance between these genome graphs will deviate further from distances between true string sets. Adding a SUD term to the objective may address this problem.

## 7 Acknowledgements

This work was supported in part by the Gordon and Betty Moore Foundation’s Data-Driven Discovery Initiative [GBMF4554 to C.K.], by the US National Institutes of Health [R01GM122935], the US National Science Foundation [DBI-1937540] and the Carnegie Mellon University School of Computer Science Sansom graduate fellowship for computational cancer research to Y.Q.

## 8 Appendix

### Procedures to generate data sets

#### 8.1 Synthetic Sets of T-cell Receptor Sequences

We construct five reference gene groups by sampling reference sequences obtained from the IMGT database [28] that represent varied diversities of V, D, J gene repertoires. The number of sequences in each group is shown in Table 2. We construct five TCR sequence groups, and each group of TCR sequences are constructed using genes from one of the five reference gene groups. To generate each TCR sequence, we randomly select a V, D and J gene from corresponding gene group, and randomly introduce *m* ∈ *{*1, 3, 5, 8, 10*}* single-nucleotide mutations to each sequence at random locations. This step is to simulate recombination and occurrences of junction single nucleotide polymorphisms (SNPs). 500 sequences are generated in each TCR sequence group.

**Table 2:** The number of unique V, J and D gene sequences in each reference gene group.

|  | Group 1 | Group 2 | Group 3 | Group 4 | Group 5 |
| --- | --- | --- | --- | --- | --- |
| TRB_V | 5 | 34 | 63 | 92 | 121 |
| TRB_J | 5 | 8 | 11 | 14 | 15 |
| TRB_D | 3 | 3 | 3 | 3 | 3 |

We construct 50 immune repertoires in five groups. Each repertoire group are constructed using simulated TCR sequences from corresponding TCR sequence group. Within each group, each sequence set contains 2–10 sequences with randomly assigned weights that sum to 100. 45 string set pairs are generated within each group.

#### 8.2 Heterogeneous sets of Hepatitis B Virus genomes

We downloaded 30 HBV genomes from each of three host species — human, bat, duck — from the NCBI virus database [29]. We construct 3 string sets for each host species. For each string set, we randomly select 5 HBV genomes from one host and randomly assign a weight to each string so that the sum of string weights in each set is equal to 100.

